# CoSIA: an R Bioconductor package for CrOss Species Investigation and Analysis

**DOI:** 10.1101/2023.04.21.537877

**Authors:** Anisha Haldar, Vishal H. Oza, Nathaniel S. DeVoss, Amanda D. Clark, Brittany N. Lasseigne

## Abstract

**Summary:** High throughput sequencing technologies have enabled cross-species comparative transcriptomic studies; however, there are numerous challenges for these studies due to biological and technical factors. We developed CoSIA (Cross-Species Investigation and Analysis), an Bioconductor R package and Shiny app that provides an alternative framework for cross-species transcriptomic comparison of non-diseased wild-type RNA sequencing gene expression data from Bgee across tissues and species (human, mouse, rat, zebrafish, fly, and nematode) through visualization of variability, diversity, and specificity metrics.

**Availability and Implementation:** https://github.com/lasseignelab/CoSIA

**Contact:** Brittany Lasseigne

**Supplementary information:** See Supplementary Files

## 1. Introduction

With the advent of high-throughput sequencing technologies (Goodwin, McPherson and McCombie 2016), there has been an explosion in the generation of gene expression data across multiple species (Mutz *et al*. 2013). Providing an excellent opportunity to leverage this available data to better study human and model organism gene expression patterns in a biomedical context, cross-species comparative studies have elucidated disease mechanisms, evolutionary patterns, and developmental differences (LoVerso and Cui 2015). However, cross-species gene expression comparison is challenging due to biological and technical factors affecting the measurements (Conesa *et al*. 2016; Chung *et al*. 2021). Previous studies have implemented a variety of comparison methods (Zhu *et al*. 2014; Sudmant, Alexis and Burge 2015; Söllner *et al*. 2017; Panahi *et al*. 2019; Wang *et al*. 2020; Bastian *et al*. 2021; García de la Torre *et al*. 2021; Liu *et al*. 2023) that involve either taking into account the evolutionary relationships between the species or rigorous statistical assumptions to account for species-level differences in gene expression (Fisher and Others 1948; Stuart *et al*. 2003; Hu, Greenwood and Beyene 2006; Lu, Rosenfeld and Bar-Joseph 2006; Lu *et al*. 2007, 2010; Campain and Yang 2010; Le, Oltvai and Bar-Joseph 2010; Tseng, Ghosh and Feingold 2012; Kristiansson *et al*. 2013). We developed CoSIA (Cross-Species Investigation and Analysis), an R package and associated Shiny app, which provides an alternative framework for cross-species RNA expression visualization and comparison across tissues and species using variability metrics. CoSIA allows users to calculate variability, diversity, and specificity metrics across *Homo sapiens* and five species commonly used in biomedical research *Mus musculus, Rattus norvegicus, Danio rerio, Drosophila melanogaster*, and *Caenorhabditis elegans*.

## 2. Implementation

CoSIA(Haldar *et al*. 2023), is a Bioconductor package accessible as part of the Bioconductor 3.17 release. CoSIA allows for relative cross-species comparison of non-diseased wild-type RNA sequencing gene expression data across tissues and species. Specifically, CoSIA implements methods for mapping gene identifiers and orthologs, visualizing gene expression by tissue and species, and comparing cross-species gene expression metrics, as shown in Fig. 1A.

**Figure 1.**
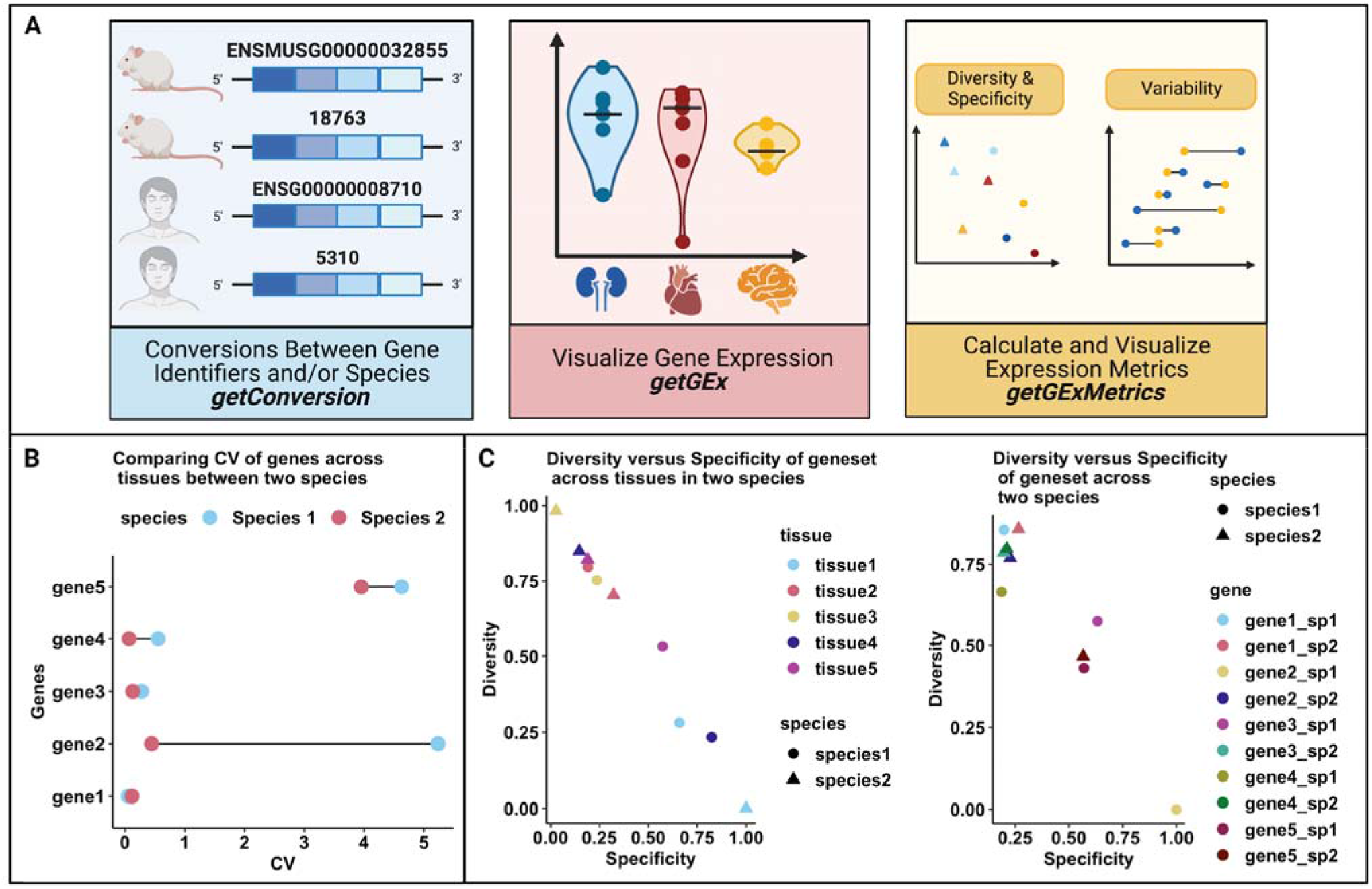
(**A**) The CoSIA R package workflow. (**B**) The coefficient of variation cross-species comparison analysis depicts a lollipop plot of example data used to show median-based CV gene comparison across species. This example highlights genes with high variability (gene5), low variability (gene4, gene3, and gene1), and species-specific variability (gene2) in expression across tissues. (**C**) The diversity and specificity cross-species comparison analysis depicts scatter plots of example data (Supplemental file: Table 2) showing diversity and specificity metric for tissues and genes. This example highlights the differences in diversity and specificity across tissues (tissue1 and tissue3 in species2), and the similarities in diversity and specificity across species (gene1, gene2, gene3, gene4 in species2). Figure was created with BioRender.com.

### 2.1 Mapping between Gene Identifiers and Orthologs (*getConversion*)

CoSIA provides the user with a streamlined method for converting between different gene identifiers and cross-species ortholog mapping. The current gene identifiers supported are Ensembl Ids, Entrez Ids, and Gene Symbols. These conversions are performed with the BiomaRt (Durinck *et al*. 2005) and AnnotationDBI (Pagès *et al*. 2022) packages, while ortholog gene mapping is performed using the NCBI Homologene Database (Sayers *et al*. 2022) and the NCBI Eukaryotic Genome Annotation Pipeline Database (Thibaud-Nissen *et al*. 2016).

### 2.2 Visualizing Gene Expression Data (*getGEx*)

CoSIA uses the Bioconductor Data Package CoSIAdata (EH7858, EH7859, EH7860, EH7861, EH7862, EH7863) hosted on ExperimentHub, which contains variance stabilized Transcript per million (TPM) gene expression values of non-diseased wild-type RNA-seq read counts we retrieved using BgeeDB [package v2.26.0; database v15.0] (Komljenovic *et al*. 2018). The variance stabilization is done using the Variance Stabilizing Transformation (VST) method implemented in the DESeq2 (Love, Huber and Anders 2014) package. These values are parsed (depending on the user’s choice of genes, tissues, and species) into a data frame that can be visualized as an interactive violin plot through the *plotSpeciesGEx* and *plotTissueGEx* plotting functions.

### 2.3 Calculating and Visualizing Gene Expression Metric Data (*getGExMetrics*)

Comparing transcriptional profiles across species has been challenging because of differences in gene expression patterns and batch effects. Previous attempts at directly comparing expression between species using various normalization techniques (Dunn *et al*. 2018) have been shown to be affected by annotation depth and quality (Oziolor, Arat and Martin 2021). Other studies (Breschi *et al*. 2016) have shown that each gene has a specific pattern, with some genes showing higher variation between organs within the same species compared to variation between species and vice versa. Direct comparison methods do not account for these aspects. To overcome these challenges, we implemented variability (Coefficient of Variation), diversity, and specificity (Shannon entropy-based) metrics that allow for relative comparison of gene expression patterns between species. To understand how these metrics work, we have simulated expression data of five genes and its orthologs across two species and five tissues (Supp. Table 2). The metrics calculated on these genes are plotted in Fig. 1B and C.

#### Coefficient of Variation (CV)

The CV in CoSIA is calculated as the standard deviation over the median using VST values. CoSIA provides two approaches for calculating the CV. ‘CV Tissue’ calculates the CV of a set of user-supplied genes across the specified tissues, while ‘CV Species’ calculates the CV of user-supplied genes across the specified species. The calculated CVs are returned as a data frame and visualized as lollipop plots using the *plotCVGEx* plotting function. As shown in Fig. 1B, Gene 2 has high variation in expression across tissues in species 1 but not in species 2. However, Gene 1 and 3 have no variation in expression across both species, thus their expression doesn’t change much across tissues in both species.

#### Diversity and Specificity

To calculate the diversity and specificity metrics, we first calculate the median of the variance stabilized TPM values for each gene in a specific tissue in a given species. These median values are rescaled using min-max scaling, which preserves the distribution of values but rescales the values between 0 and 1. These values are used to calculate the diversity and specificity metrics as described in (Martínez and Reyes-Valdés 2008). Briefly, diversity and specificity in the context of gene expression across tissues is quantified using Shanon entropy. Diversity refers to the degree of heterogeneity or variability in the expression patterns of a gene across different tissues. It captures the distribution and relative frequency of expression across tissues, with higher diversity indicating a more evenly distributed expression pattern. Specificity, on the other hand, measures the level of concentration or selectivity of gene expression within a particular tissue. It assesses the extent to which a gene’s expression is confined to a specific tissue, with higher specificity indicating a more restricted or specialized expression pattern within that tissue. The range of diversity and specificity is between 0 and 1, with 0 being low and 1 being high. Another important thing to note is that they are inversely related (Jones *et al*. 2023).. Thus, a highly diverse gene will have similar expression across tissues but low specificity. A highly specific gene will have higher expression in one tissue compared to other tissues. CoSIA allows for four different calculations of diversity and specificity in a species: i) ‘DS Gene’ compares user-specified genes across user-specified tissues, ii) ‘DS Gene all’ compares user-specified genes across all tissues in a species, iii) ‘DS Tissue’ compares user-specified tissues across user-specified genes, and iv) ‘DS Tissue all’ compares user-specified tissues across all genes in a species. The metrics can either be exported as a data frame or visualized using diversity/specificity scatter plots using the *plotDSGEx* plotting function. The first half of Fig. 1C, shows that the geneset expression is very specific in tissue 1 in species 2 compared to species 1. On the contrary, the geneset expression is very diverse in tissue 3 in species 2 compared to species 1. In the second half of Fig 1C, we look at individual genes, where gene 2 has very specific expression in species 1 however, it is very diverse in species 2.

### 2.4 Shiny App

We have also implemented the CoSIA package as a Shiny app (DeVoss et al. 2023) hosted at (https://lasseignelab.shinyapps.io/CoSIA/) which provides a graphical user interface (GUI) with similar functionalities as the package.

## 3. Conclusion

Direct comparison of gene expression between species is complex as it is confounded by differing levels of expression and function of orthologous genes across species. Here, we provide the CoSIA package and Shiny app to facilitate the relative comparison of gene expression by summary metrics (i.e., coefficient of variation, diversity, and specificity) in six species. By leveraging the Bgee database of species-specific RNA-Seq expression data, we provide tools for the robust comparison of gene expression values across both species and tissues. We believe CoSIA will also aid biomedical researchers in selecting optimal model organisms for a given gene in a tissue of interest.

## Acknowledgements

The authors would like to thank the members of the Lassigne lab for their support and feedback, in particular, Elizabeth J. Wilk, Jordan Whitlock, and Timothy C. Howton.

## Supplemental Files

CoSIA_AppNote_Supplementary_Data.docx contains the following: (1) *Table 1* - Table of Package Dependencies for CoSIA, (2) *Table 2*- Median Variance Stabilized Transformation of RNA-seq Read Counts used to render Figure 1b and 1c (3) Figure S1 - CoSIA Usage Example: cross-species & -tissue investigation of genes associated with monogenic kidney diseases.

