## Supplemental for "CoSIA: an R Bioconductor package for CrOss Species Investigation and Analysis"

**Table 1.** Package Dependencies for CoSIA. This table includes the package name and minimum version of the package required for use in CoSIA. Entries are sorted alphabetically.

| **Package Names** | **Minimum Version Required for CoSIA** |
| --- | --- |
| AnnotationDbi | 1.52.0 |
| annotationTools | 1.64.0 |
| BgeeDB | 2.26.0 |
| BiocStyle | 2.22.0 |
| biomaRt | 2.46.3 |
| dplyr | 1.0.7 |
| ExperimentHub | 2.7.0 |
| ggplot2 | 3.3.5 |
| homologene | 1.4.68.19 |
| knitr | 1.42 |
| magrittr | 2.0.1 |
| methods | 4.1.2 |
| org.Ce.eg.db | 3.12.0 |
| org.Dm.eg.db | 3.12.0 |
| org.Dr.eg.db | 3.12.0 |
| org.Hs.eg.db | 3.12.0 |
| org.Mm.eg.db | 3.12.0 |
| org.Rn.eg.db | 3.12.0 |
| plotly | 4.10.0 |
| qpdf | 1.3.0 |
| RColorBrewer | 1.1-2 |
| readr | 2.1.1 |
| rmarkdown | 2.20 |
| stats | 4.1.2 |
| stringr | 1.4.0 |
| testthat | 3.1.6 |
| tibble | 3.1.7 |
| tidyr | 1.2.0 |
| tidyselect | 1.1.2 |
| tidyverse | 1.3.1 |

**Table 2.** Median Variance Stabilized Transformation of RNA-seq Read Counts example data used to render Figure 1B and 1C

|  | tissue1 | tissue2 | tissue3 | tissue4 | tissue5 | species |
| --- | --- | --- | --- | --- | --- | --- |
| gene1_sp1 | 8.34 | 8.55 | 8.65 | 7.56 | 8.55 | species1 |
| gene2_sp1 | 1.6 | 1.6 | 1.6 | 1.6 | 20.34 | species1 |
| gene3_sp1 | 5.43 | 6.34 | 5.86 | 9.4 | 5.83 | species1 |
| gene4_sp1 | 6.34 | 12.35 | 12.35 | 6.32 | 5.32 | species1 |
| gene5_sp1 | 2.3 | 21.34 | 22.12 | 2.3 | 2.3 | species1 |
| gene1_sp2 | 4.54 | 3.45 | 4.67 | 4.78 | 4.66 | species2 |
| gene2_sp2 | 4.5 | 6.7 | 12.4 | 12.7 | 18.43 | species2 |
| gene3_sp2 | 1.3 | 1.4 | 1.6 | 1.8 | 1.7 | species2 |
| gene4_sp2 | 7.3 | 8.4 | 8.6 | 8.4 | 7.6 | species2 |
| gene5_sp2 | 1.8 | 18.2 | 17.4 | 1.8 | 2.2 | species2 |

**Figure S1.**

**A)**	**B)**

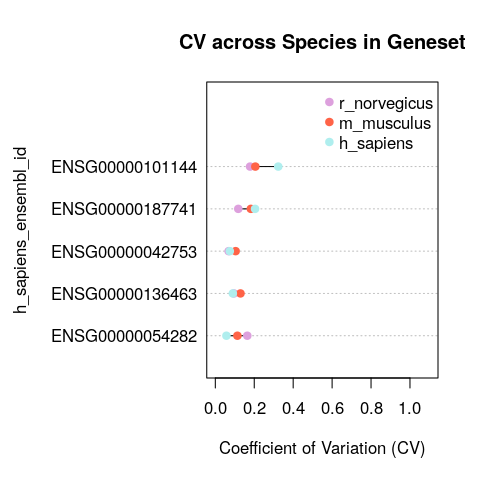

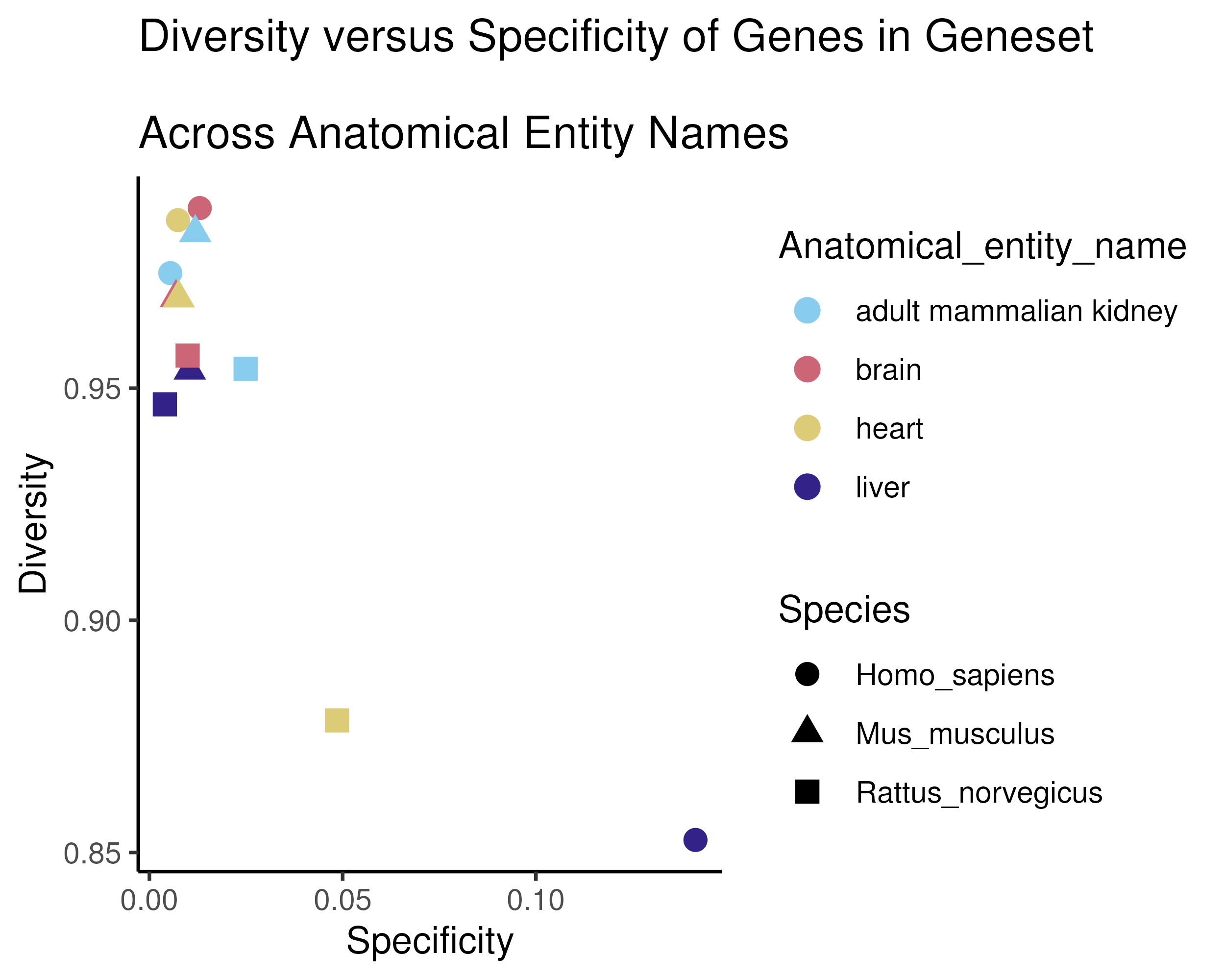

CoSIA metrics calculated on genes associated with monogenic kidney diseases in humans and its orthologs in mice and rats **(A)** The coefficient of variation cross-species comparison for **(B)** Diversity and specificity metrics across kidney, brain, heart, and liver.
